# Generation and Culture of Cardiac Microtissues in a Microfluidic Chip with a Reversible Open Top Enables Electrical Pacing, Dynamic Drug Dosing and Endothelial Cell Co-Culture

**DOI:** 10.1101/2021.11.01.465885

**Authors:** Aisen Vivas, Camilo IJspeert, Jesper Yue Pan, Kim Vermeul, Albert van den Berg, Robert Passier, Stephan Sylvest Keller, Andries D. van der Meer

## Abstract

Cardiovascular disease morbidity has increased worldwide in recent years while drug development has been affected by failures in clinical trials and lack of physiologically relevant models. Organs-on-chips and human pluripotent stem cell technologies aid to overcome some of the limitations in cardiac *in vitro* models. Here, a bi-compartmental, monolithic heart-on-chip device that facilitates porous membrane integration in a single fabrication step is presented. Moreover, the device includes open-top compartments that allow facile co-culture of human pluripotent stem cell-derived cardiomyocytes and human adult cardiac fibroblast into geometrically defined cardiac microtissues. The device can be reversibly closed with a glass seal or a lid with fully customized 3D-printed pyrolytic carbon electrodes allowing electrical stimulation of cardiac microtissues. A subjacent microfluidic channel allowed localized and dynamic drug administration to the cardiac microtissues, as demonstrated by a chronotropic response to isoprenaline. Moreover, the microfluidic channel could also be populated with human induced pluripotent stem-derived endothelial cells allowing co-culture of heterotypic cardiac cells in one device. Overall, this study demonstrates a unique heart-on-chip model that systematically integrates the structure and electromechanical microenvironment of cardiac tissues in a device that enables active perfusion and dynamic drug dosing. Advances in the engineering of human heart-on-chip models represent an important step towards making organ-on-a-chip technology a routine aspect of preclinical cardiac drug development.

## 1. Introduction

Cardiovascular diseases (CVD) are the leading cause of death in western society. In Europe alone, nearly 4 million people die every year due to CVD. ^[1]^ Drug development for CVD has been hindered, among others, by reduced investments in drug research due to multiple failed late-phase clinical trials. ^[2]^

After years of development, drugs can fail in human trials for a variety of reasons, e.g., lack of efficacy or toxicity. Such unexpected outcomes of clinical trials are a clear illustration of the low correlation between results in common cell culture models and animal models on the one hand, and human subjects on the other. ^[3]^ This low correlation often stems from physiological and metabolic differences between animal models and humans. ^[4]^

With the advent of human stem cell technology, some of these hindrances can now be overcome. Human stem cells can be differentiated into tissue-specific cells that can be used for pre-clinical testing in drug development. Successful examples include the use of stem cell-derived cardiomyocytes for cardiotoxicity testing. ^[5]^ Notwithstanding, cells derived from human pluripotent stem cells (hPSC), such as human embryonic stem cells (hESC) or human induced pluripotent stem cells (hiPSC), are often in an immature state and require more complex culture systems than basic 2D well plates to promote their maturation or organization into functional tissues.

Traditional culture well plates do not provide the ideal microenvironment for stem cell cultures, lacking complex cell-cell and cell-extracellular matrix (ECM) interactions that have been recognized to play crucial roles in cell maturation and drug responses. ^[6]^ Moreover, essential biophysical aspects are missing in these conventional cell culture platforms such as geometrical cues to enhance muscle fiber alignment in hPSCs-derived cardiomyocytes (hPSC-CM) or shear stress stimulation on hPSCs-derived endothelial cells (hPSC-ECs), both of which are important for e.g. metabolic responses and recapitulation of the human cell microenvironment. ^[7]^

Various approaches to mimic relevant aspects of the microenvironment have successfully been implemented in cardiac *in vitro* models to improve their functional relevance ^[8–10]^. Using microfabrication techniques, multielectrode arrays (MEAs) have been used to provide electrical sensing and stimulation to cultured cardiomyocyte monolayers. ^[11]^ Mechanical stimulation has been achieved in 2D using cultures on flexible muscular thin films that allow non-invasive sensing of contractile force and beating frequency. ^[12]^ Integration of multiple cardiac cell types in 3D spheroids has been shown to increase the resemblance with the cardiac microenvironment in healthy and diseased states. ^[13,14]^ Finally, by coupling co-cultures with electromechanical stimulation, engineered heart tissues (EHTs) and cardiac wires have been able to mimic relevant key aspects of cardiac physiology, such as heterotypic cell interaction, electrical actuation and biochemical cues that altogether capture many aspects of cardiac physiology and have been shown to enhance hPSC-CM maturity. ^[15,16]^

Organs-on-chips (OoCs) are microfluidic cell culture devices that allow culture of cells in dynamic conditions with tight control over the physico-chemical microenvironment. ^[17,18]^ Moreover, OoCs allow the integration of sensors to obtain *in situ* readouts. ^[19,20]^ Many OoC systems that recapitulate certain aspects of cardiac function, known as ‘hearts-on-chips’ (HoC), have been reported. ^[21,22]^ Integration of the advances in cardiac cell culture with HoC technology has been done on a ‘fit-for-purpose’ basis, where designs are guided by one specific aspect of cardiac physiology. Examples of this are HoCs with relatively simple mono-cultures that focus on electrophysiology ^[12,23,24]^ or on geometrical confinement. ^[25,26]^ Others have focused on more complex heterotypic co-cultures to realistically capture the cardiac tissue architecture. ^[16]^ Reported HoC devices with such complex co-cultures either have a planar configuration with side-by-side microfluidic compartments, ^[27,28]^ or they have a complex 3D configuration with advanced perfusable scaffolds. ^[29–31]^

One of the many advantages of OoCs is the ability to recreate *in vitro* tissue interfaces by engineering two stacked microcompartments separated by a semi-permeable membrane. Additionally, the increasing evidence of the role that the endothelium plays in cardiac physiology requires more representative models to study these mechanisms. However, no HoC has been reported that makes use of 3D cardiac tissue culture in combination with a perfusable microchannel in such a typical multi-layer configuration. This configuration would offer unique advantages, such as control on the relative position, localized drug dosing and proximity of cardiac microtissues and an endothelial monolayer, ease of fabrication and selective perfusion of compounds in separate compartments. Combining cardiac and vascular cells in this configuration would help to establish models that better mimic the human cardiac anatomy. Increasing awareness of the role that the endothelium plays in heart function by the cross-talk between these cells, such as the nitric oxide effects and cytokine release upon infection, also support the creation of more complex *in vitro* models that are able to recapitulate some aspects of heart physiology. It would also allow seamless future integration of such HoCs into multi-organ ‘body-on-a-chip’ systems, which rely on linking organs-on-chips via microchannels.

Here, we report the development of a new versatile HoC device that mimics 3D cardiac tissue geometry by employing a unique multi-layer organ-on-chip design with a reversible open-top configuration and an easy one-step fabrication process resulting in a monolithic device. The open top enables direct seeding of hPSC-CMs and cardiac fibroblasts in microwells with defined 3D geometries in close proximity with a subjacent microfluidic channel where a drug can be locally administered and be lined by hPSC-ECs. The open top also allows reversible integration of 3D-printed carbon electrodes for stimulation of the cardiac tissues. Having a multi-layer configuration allows selective perfusion of each compartment with a defined medium of choice. Moreover, we use the HoC to demonstrate that highly localized and dynamic drug dosing of cardioactive drugs through the microfluidic channel more closely mimics the kinetics of drugs in cardiac tissue

## 2. Results and discussion

### 2.1. Heart-on-Chip Design and Fabrication

The HoC device was designed to establish a controlled 3D co-culture with cardiac tissues and endothelial cells in a perfusable device. The compartmentalization of the device and the membrane dividing both compartments made it possible to fluidically stimulate the endothelial cells without disturbing the cardiac tissues.

As depicted in **Figure 1**A, the HoC hosts eight cardiac microtissues in two fluidically independent culture chambers. In Figure 1B left a top schematic view of the HoC is shown demonstrating the integrated porous polyester membrane and the monolithic design of the HoC.

**Figure 1.**
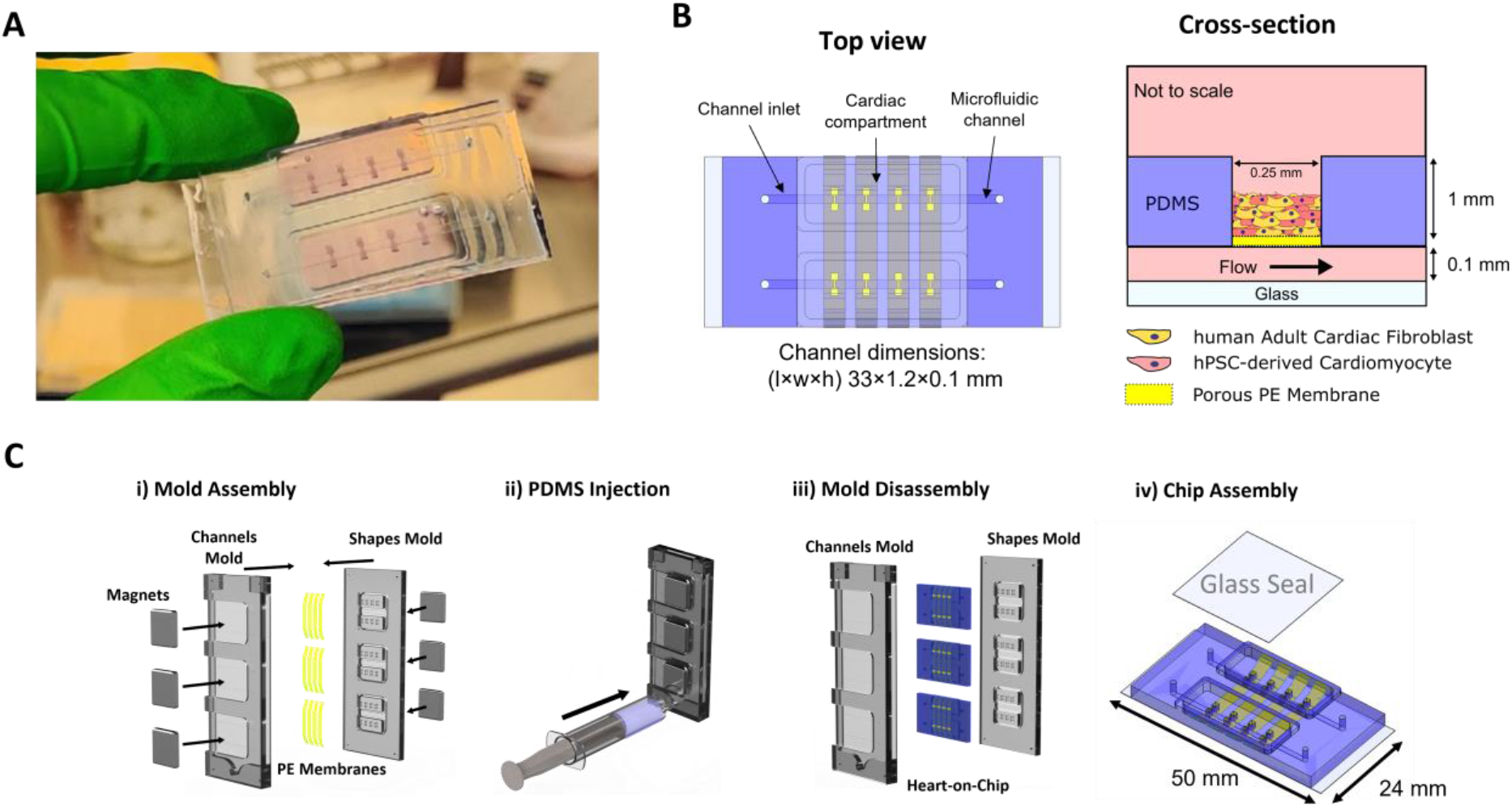
Heart-on-chip features, configuration and fabrication process. A) Picture of the heart-on-chip device. B) on the left, the top schematic view of the heart on chip device depicting the monolithically integrated porous membrane in yellow; on the right, schematic cross-sectional view of the culture compartments (note: not to scale), the cardiac microtissues, its respective cell types and its real dimensions. C) General overview of the fabrication process of the heart-on-chip. i) Membranes are cut and fixed on the shapes mold using double sided tape; the mold parts were then brought together and the magnets applied in their respective pockets ensuring application of uniform pressure. ii) PDMS is degassed and injected into the mold using a syringe, bubbles are allowed to escape the mold and cured in an oven; iii) after PDMS curing, molds were separated and the heart-on-chip devices were diced after being peeled of the mold; iv) isometric schematic view of the assembled heart-on-chip device and the glass seal used to reversibly close the cardiac compartment and respective dimensions.

Similar to compartmental pharmacokinetics assays, ^[32]^ where each organ is represented by a compartment, the HoC design takes this philosophy into consideration by creating two compartments depicted in Figure 1B right that can be selectively perfused with a medium of choice. The shape of the 3D cardiac tissue compartment was adopted from earlier studies ^[25]^ that reported on the benefits of using such a geometry for the uniaxial alignment of the fabricated cardiac microtissues’ muscle fibers and it could potentially be altered to another geometry of choice. Cytoskeletal organization is considered a hallmark of cardiac tissue functionality and contractility. Therefore, promoting this configuration enhances the recapitulation of the organ structure in the HoC. ^[33]^

The cardiac microtissues were reversibly enclosed in the open-top compartments in the top layer of the device. This open-top configuration enables facile seeding of cardiac cell suspensions, medium sampling and cardiac microtissue retrieval at any point of the culture and prevents detrimental media evaporation. Moreover, when reversibly sealing the top compartment with a glass seal, perfusion of the microchannel could be performed with flow rates and pressures of up to 147 μL/min and 18 mbar. The sealing of the glass seal relied on evaporation and subsequent formation of protein deposits in a pre-designed micropatterned zone around the edge of the full top compartment without any glue or tape (**Supplementary Figure 1**). When double-sided tape was applied, the seal strength could be further improved, allowing flow rates and positive pressures of up to 339 μL/min and 65 mbar, respectively.

The ability to selectively perfuse the microfluidic channel allows perfusion of controlled concentrations of drugs through the microchannel for localized administration of drugs to the cardiac microtissues. This is a relevant feature for drug studies, where localized drug administration is extremely useful to gain insight into how cells in a tissue react to a specific drug at different concentrations.

The fabrication of the device was designed to be simple, requiring as few steps as possible employing an injection-molding-like fabrication technique adopted from. ^[34]^ This method allows fabrication of multiple HoCs in a one-step process, yielding monolithic devices with integrated porous membranes that do not require the use of toxic glues or multi-step protocols typically needed to integrate porous membranes in multicompartment organs-on-chips (Figure 1C, i). ^[35]^ Additionally, this method prevented issues with fluid leakage and infections that could occur due to the open conduit the medium typically forms with the surrounding space. The neodymium magnets employed in the chip fabrication were able to provide an evenly distributed and consistent clamping force that prevented the PDMS infiltration over the porous membrane, which would otherwise obstruct the fluid link between the two culture compartments. Recently, we demonstrated that the same technique also allows facile integration of other membranes in organ-on-chip devices, including biological membranes of varying thicknesses. ^[36]^

Injection of PDMS into the molds generated bubbles that naturally escaped via the top opening of the molds after room temperature incubation, leaving the mold bubble free (Figure 1C, ii). After the PDMS curing step, the removal and dicing of the HoCs was performed with caution to not separate the integrated membranes from the PDMS (Figure 1C, iii). Finally, the slabs of PDMS were bonded to a glass coverslip with spin-coated PDMS. The PDMS on the coverslip provided a homogeneous environment to the cells in the microchannel and practically prevented bonding issues between the coverslip and the monolithic HoC (Figure 1C, iv).

### 2.2. Cardiac microtissues in the Heart-on-Chip

The role that the microenvironment plays in cell culture highlights the relevance of integrating 3D cultures in HoC devices. Significant differences have been found between 2D and 3D cardiac cultures with respect to their maturity and functionality. ^[14]^ Moreover, the integration of multiple cardiac cell types in 3D cardiac cultures has been shown to promote maturity markers in CMs and better recapitulate *in vitro* the microenvironment these cells would experience *in vivo*. ^[13]^ These characteristics highlight the advantages that cardiac microtissues have over 2D cultures.

We integrated cardiac microtissues that were fabricated using a human embryonic stem cell line with two reporter genes for the transcription factor NKX-2.5^GFP/+^ and the fluorescently tagged sarcomere accessory protein α-actinin with GFP and mRuby, respectively. ^[37]^ NKX-2.5 is a cardiac-specific marker used to identify hPSC committed to a cardiac phenotype ^[38]^ while α-actinin allows the visualization and assessment of the contractile units of these cells. Ensuring uniform cardiac microtissue fabrication is crucial to guarantee reproducibility of drug exposure assays since differences in sizes may alter the dynamics of tissue oxygenation and drug concentration seen by the cells. To that end, we optimized the fabrication of cardiac microtissues by working with well-characterized batches of cells and by identifying the optimal ratio of CMs and cFBs. We characterized the percentage of CMs present in the cell population obtained after differentiation using flow cytometry and the reporter genes of the hPSC-CMs (**Supplementary figure 2**). The percentage of NKX-2.5 and α-actinin positive cells was 67 ± 11% (average ± STD). Even though the differentiation efficiency was relatively consistent, introduction of an additional cardiac cell type, human adult cFB, at a ratio of CMs/cFB of 1:10 improved the cardiac tissue compaction (**Supplementary figure 3**). cFBs highly improved the consistency of the cardiac microtissues by allowing the tissues to remodel and compact inwards and pinning them in the shaft of the cardiac compartment. By applying these optimized cell culture conditions, we were able to obtain homogenous compaction among the fabricated tissues in a consistent manner after three days of culture (**Figure 2**A). Moreover, cardiac microtissues in culture kept beating for a maximum period of 30 days with three changes of medium per week. The extended culture period may be important for future applications, as it allows cells in the tissue to form heterotypic and homotypic cell connections for their electromechanical coupling and paracrine communication. ^[39]^

**Figure 2.**
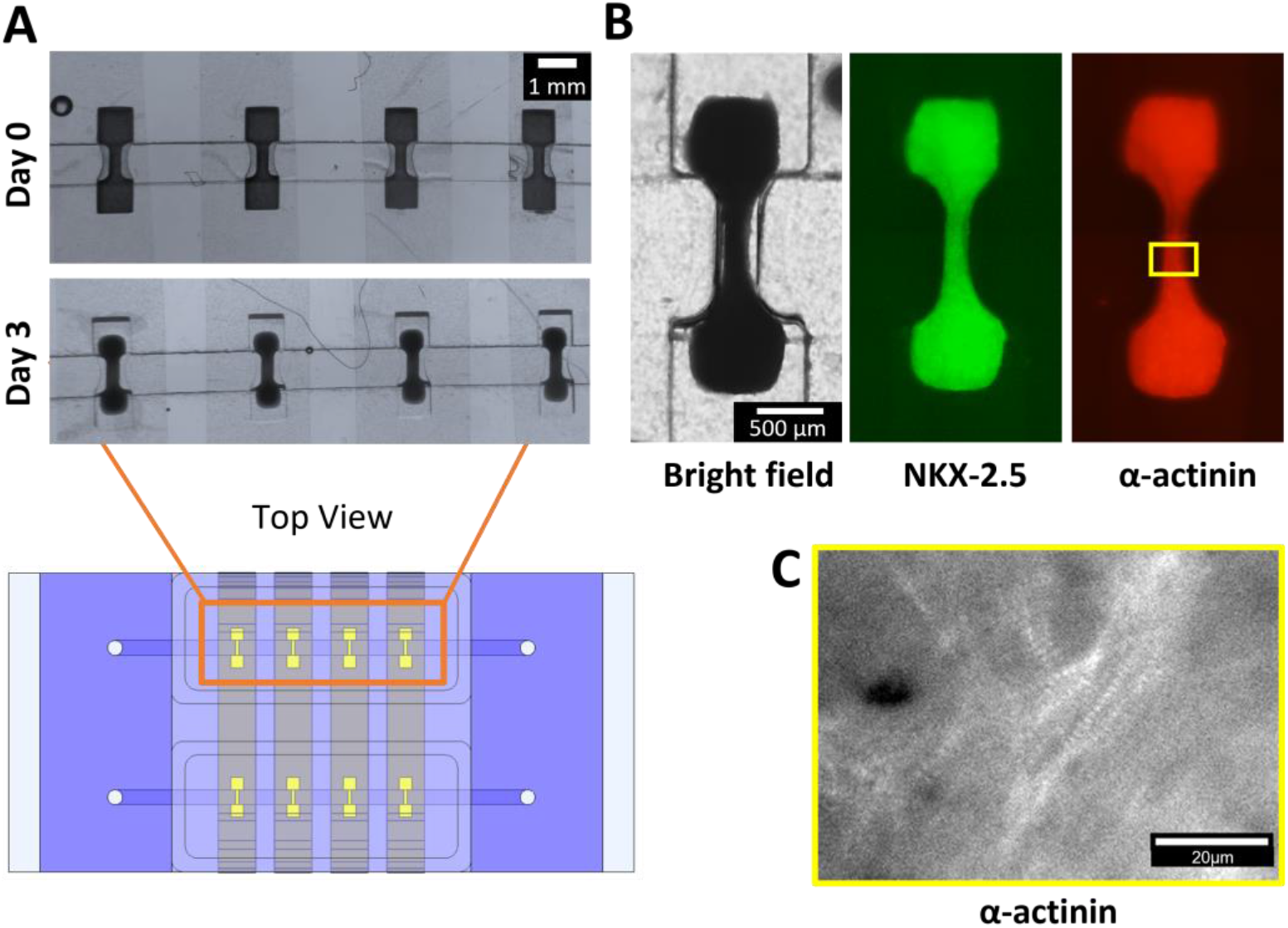
Cell culture in the cardiac and microchannel of the heart-on-chip. A) On day 0 the hydrogel-cell mix fills the entire dumbbell-shaped compartment and after 3 days the tissues have compacted and pinned in the shaft of the compartment. B) Close-up on one 3D cardiac tissue in bright field and fluorescent microscopy using the GFP reporter gene NKX-2.5 and the fluorescently tagged α-actinin sarcomeric protein. C) Close-up at the single cell level showing the striated sarcomeric structures of hPSC-CMs in the 3D cardiac tissue using the fluorescently tagged α a-actinin protein. Scale bar 20 μm.

The NKX-2.5 reporter gene in the hPSC-CMs proved useful to determine cell distribution in the cardiac microtissues without laborious staining protocols. Based on fluorescence microscopy, the CMs were evenly distributed throughout the cardiac microtissues (Figure 2B). This homogeneity also indicated that there were no cFBs clusters in the formed tissues. In case a non-homogenous distribution of cFBs would have been observed, this could have resulted in the formation of cardiac microtissues resembling a fibrotic state. ^[16]^

In turn, fluorescently tagged α-actinin allowed analysis of the organization of the sarcomeres in live samples and throughout an experiment (Figure 2B). In the present study, we were able to verify the striated pattern characteristic of the sarcomeric structure of the CMs due to the α-actinin/mRuby fusion protein. The presence of these structures indicates the maturity of the contractile machinery of CMs and even though not studied here, it could enable the assessment of the sarcomeric structural mutations involved in dilated cardiomyopathy. ^[40]^

In this study, no obvious sarcomeric alignment was seen in the cardiac microtissues (Figure 2C). It has been reported that cardiac tissues require an anchorage point in order to increase the uniaxial contraction. ^[25]^ In our case, the tissues were not anchored to the PDMS substrate of the squared-shaped region of the dumbbell-shaped cardiac compartment but pinned in the shaft of said compartment. Even after chemical functionalization to promote anchorage of CMs to PDMS, ^[41]^ tissues did not anchor to the bottom of the squares of the cardiac compartment, which may be the reason for the relatively randomly oriented sarcomeric structures.

Overall, a protocol for cardiac microtissue fabrication in a systematic and standardized manner was established and successfully characterized. In order to reduce the variability of the CMs population inherent to hPSC-CM differentiation, an extra step of purification should be implemented. Alternatives to do so include cell sorting techniques and chemical purification using nutrient selective protocols that promote cardiac cell viability while reducing non-cardiac cells viability by lactate as an energy source. ^[42]^

### 2.3. Electrical stimulation of cardiac microtissues using an E-lid with 3D-printed pyrolyzed carbon electrodes

With each heartbeat, cardiac cells synchronously contract after being triggered by an action potential generated by the sinoatrial node – the heart pacemaker. To mimic this mechanism, depolarization of CMs using electrical pulses has been used *in vitro*. Thus, electrical pacing is a valuable feature to include in HoCs devices and reviewed elsewhere in more detail. ^[43]^ Integration of electrical pacing aids in the maturation of CMs further mimics the *in vivo* microenvironment. ^[15]^

We were able to pace the cardiac microtissues in our HoC using an electronic lid (‘E-lid’) comprised of highly customizable 3D-printed pyrolytic carbon electrodes (**Figure 3**A and B) instead of the glass seal (Figure 1C iv). The E-lid encompasses a set of eight pairs of pyrolytic carbon micropillars electrodes with a height of 1.45±0.16 mm and a diameter of 158±8 μm connected to two electrical carbon pads that allow the connection to a signal generator of choice (Figure 3A). Designs with different dimensions were tested in order to determine the dimensions of the micropillars electrodes that would best fit in the dumbbell-shaped wells of the cardiac compartment. The effects of printing accuracy and mass loss during the pyrolysis process of the micropillar electrodes can be appreciated in **Supplementary Figure 4**. Due to the high conductivity of carbon and its low secondary reactions with the culture medium, ^[44]^ common hurdles in long-term electrical stimulation assays with metallic electrodes can be overcome. More importantly, carbon biocompatibility allowed seamless integration of the electrodes and cell attachment to the electrodes. During pacing, the electrodes did not show any signs of drastic disturbances of the cell culture microenvironment, medium pH changes or bubbling. Despite the good attachment of the cells to the pillar electrodes, no obvious alignment could be detected by using a fluorescence inverted microscope and the fluorescently tagged reporter α-actinin gene. Two-photon microscopy or transmission electron microscopy are techniques that can be employed to better elucidate the sarcomeric structural organization of the tissues in these conditions.

**Figure 3.**
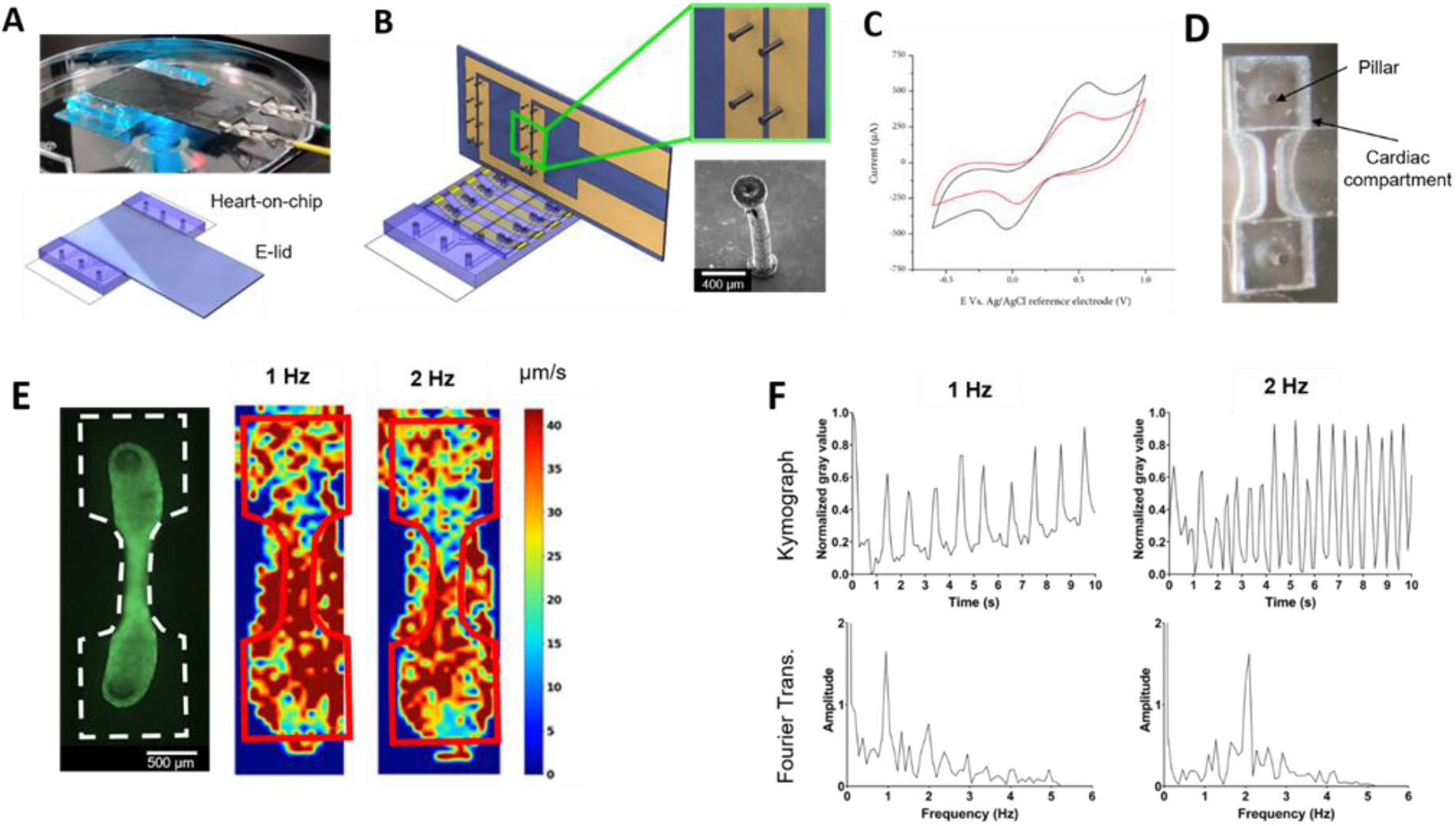
E-lid with carbon electrodes integrated in the heart-on-chip. A) The glass seal is replaced by the E-lid shown in a schematic form, bottom, and in a picture depicting the connection of the E-lid to the stimulus generator - top. B) Schematic view of the opened device depicting the HoC and the E-lid and the respective electrodes. A schematic view of the electrodes is shown in the top right and in the bottom right a scanning electron microscopy image shows one of the 3D printed pyrolytic carbon electrodes with a height of 1.45 mm and a diameter of 158 μm. C) Cyclic voltammogram recorded with 10 mM Ferri/ferrocyanide at a scan rate of 50 mV/s comparing an E-lid with flat carbon electrodes (red) with one having 3D pillars (black). D) Alignment of the electrodes in the dumbbell-shaped compartment without cells. E) Fluorescent picture of a 3D cardiac tissue attached to the carbon electrodes in monoculture on the left-hand side and in the center and on the right a motion heatmap of the cardiac tissues generated with optical flow image analysis using OpenHeartWare script. F) Beating frequency plots of a cardiac tissue obtained using kymography and their respective Fourier transform evidencing the dominant pacing frequency in the cardiac tissue beating pattern at 1 Hz and 2Hz.

We measured the electrical conductivity of the carbon leads with the electrodes, which was 80 ± 13 mΩcm, (average ± STD), which is similar to what has been shown before for 2D pyrolytic carbon. ^[45]^ Additionally, we also employed cyclic voltammetry to verify if the 3D-printed pyrolytic carbon electrodes were conductive and had an electroactive surface able to provide electrical stimulus to the cardiac microtissues (Figure 3C). The considerably higher anodic and cathodic peak currents for the 3D carbon electrodes confirmed an increased electrode surface area compared to 2D electrodes without the 3D pillars. This demonstrated a successful structural and electrical integration of highly conductive 3D carbon pillars on the E-lid required for *in situ* stimulation of the cardiac microtissues. Furthermore, the increased surface area for signal transduction and close interaction with the cardiac microtissues and the E-lid’s electrodes allows reducing the applied potential difference between the electrode pairs to elicit an action potential. This reduces undesirable water dissociation and culture medium acidification typically seen in 2D or Pt wire electrodes, which is detrimental in electrical stimulation experiments. Further characterization of the fabricated electrodes using electrical impedance spectroscopy would be required to better understand the electrochemical phenomena taking place during stimulation with these electrodes, as well as the characterization of their charge storage capacity which is a relevant parameter in electrode stimulation design.

After injection of the cardiac cell suspensions, but prior to hydrogel gelation, the E-lid was used to seal the device using double-sided tape. The electrodes were centered in the cardiac compartment ensuring equal distribution of cells around them (Figure 3D). Cardiac microtissues were cultured for at least 10 days prior to electrical stimulation. During this period, cells compacted and wrapped around the carbon electrodes and were autonomously beating, due to the biocompatibility of carbon as an electrode material and an increase in electrode-tissue contact area (Figure 3E, left).

Exposure of cells to electrical fields leads to a charge imbalance in the extracellular space created by the ion migration towards the electrodes of opposing charge. Due to the relatively high impedance of the cellular membrane, the cytosolic ion distribution remains unaltered. The anisotropic charge distribution in the extracellular and intracellular space hyperpolarizes one extremity of the cells while depolarizing the other. When a threshold is reached, the action potential is triggered and thus cell contraction takes place. ^[46]^ In cardiac tissues, however, the description of this phenomenon is more complex due to the heterogeneity of the tissue and the presence of cellular gap junctions that electrically couple the cells that populate the tissue and alter signal propagation. The details of this phenomenon has been reviewed elsewhere in more depth. ^[46]^ Briefly, cells near the electrodes depolarize and trigger an action potential that propagates along the tissue due to the presence of gap junctions in the cells.

In order to determine the excitation threshold (ET), i.e., the minimum potential difference required to elicit a tissue contraction, we progressively increased the potential difference between the electrodes from 10 mV up to 350 mV. Typically, the ET is visually determined and subsequent stimulus is performed as high as 200% the ET to ensure that all cells are being effectively stimulated and with common values ranging from 2 to 10 V. ^[15,47,48]^ Increasing the potential difference may introduce undesirable effects such as hydrolysis in the culture medium. Moreover, visual detection of the ET is a subjective measure that may introduce errors due to user bias. To overcome this limitation, we used the Fourier transform of the beating signal to determine the ET. Thus, the ET was established as the minimum potential difference where the electric stimulation signal frequency was identified in the Fourier transform of the tissue beating signal. Using this method, the ET was determined to be at 350 mV (**Supplementary Figure 5**).

At this potential, cardiac tissues were exposed to an estimated electrical field of 1.6 V/cm, sufficient to achieve synchronized beating at 1 Hz and 2 Hz (Figure 3F). This field strength is below the minimum 2 V/cm value reported in literature. ^[15,16]^ Due to the proximity and increased contact area between the CMs in the cardiac tissue and the electrodes, we were able to reduce the electric field required to capture the cardiac tissues contraction, and consequently reduce water hydrolysis when compared to platinum wires electrodes. Avoiding these electrochemical reactions allows to increase pacing efficacy, electrode life and to avoid disturbances of the cell microenvironment. The optional integration of the E-lid further increases the versatility of the HoC providing integrated electrical stimulation over longer periods of time.

### 2.4. Localized drug dosing to the heart-on-chip

To test the ability of localized drug administration and the functional aspects of the formed cardiac microtissues in the HoC device, we stimulated them with isoprenaline. Isoprenaline is a non-selective β-adrenoreceptor agonist and has similar effects as epinephrine. The effect of isoprenaline on cardiac cells is relatively well-characterized. It is known to have a positive chronotropic effect, i.e. increase the beating frequency. ^[49]^

The drug was dynamically administered through the subjacent microfluidic channel, allowing for a localized drug administration to the cardiac microtissues through the porous membrane that allows fluidic communication with the cardiac compartment. Additionally, the ability to perfuse the bottom compartment allows an unperturbed microenvironment of the top cardiac compartment where the cardiac microtissues are cultured, reducing sudden microenvironmental changes that occur when spiked medium is exchanged in the device. This feature of the HoC is also found in other HoC devices in literature. ^[50,51]^ However, the devices reported so far do not integrate full 3D cardiac microtissues.

To test the localized dynamic drug administration in the HoC, the cardiac microtissues were cultured for a period of 14 days prior to the assay. On day 14, the beating rate of the cardiac microtissues was recorded and served as a baseline. Isoprenaline was then administered in spiked medium through the subjacent channel, rocked and incubated for 5 min inside an incubator with 5% CO_2_ and at 37°C (**Figure 4**A, B left). The process was repeated with increasing spiked medium concentrations - ranging from 1 to 30 nM. Similar *in vivo* plasma concentrations of 0.39 ng/mL (1.85 nM) after isoprenaline intravenous administration have been previously reported. ^[52]^

**Figure 4.**
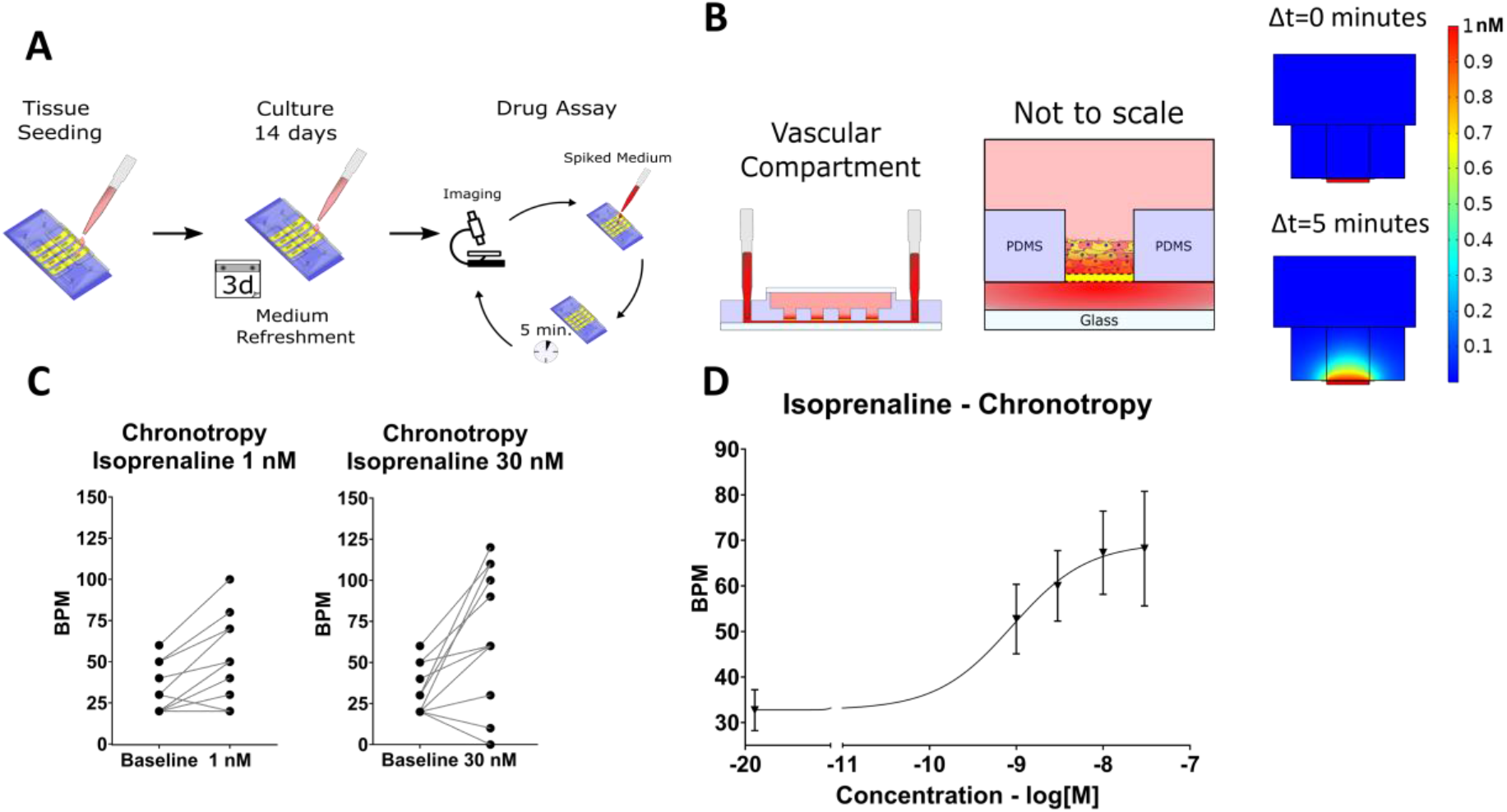
General overview of the drug dosing in the heart-on-chip, CFD diffusion study and chronotropic response of the cardiac microtissues. A) Cardiac tissues were seeded and cultured for 14 days prior to drug administration with medium changes every 3 days. A baseline of the beating frequency was recorded, drug was administered and incubated for 5 minutes and the new beating frequency was recorded. B) Localized drug administration was performed through the subjacent microfluidic channel. A cross-sectional schematic view illustrates the drug administration path (left) and the zoomed in view of the drug diffusion from the microfluidic channel into the cardiac compartment (middle). In the CFD analysis the drug diffusion at 1 nM during a 5 minute incubation was studied for the incubation period (right). C) Overall increasing trend of the individual cardiac microtissues chronotropic response for the lowest and highest isoprenaline concentration locally administered through the microfluidic channel. D) An overall increasing trend was detected in all tested concentrations of isoprenaline for the different tested concentrations.

To gain insight into the amount of drug reaching the cardiac tissues when administered via the subjacent channel after a 5 min incubation period, computational fluid dynamic (CFD) and mass transport modelling of the drug diffusion through the porous membrane was performed at the lowest concentration administered, 1 nM. The CFD analysis revealed that after the 5 minute incubation period, the cardiac microtissues would be exposed to a concentration equal to that of the spiked medium (Figure 4B right). Hence, the localized administration of the drug through the subjacent channel was able to expose the cardiac tissues to the intended drug concentration without disturbing the cardiac compartment microenvironment.

In terms of biological response, an increasing trend in the beating frequency of the individual cardiac microtissues was observed for most of the tested cardiac microtissues at any of the given tested concentrations. Figure 4C depicts the chronotropic response of each of the tissues tested at the lowest and highest concentration. A summary graph of all the tested concentrations is available in **Supplementary figure 6**. Overall, an increasing beating frequency trend upon administration of isoprenaline could be observed, with a clear dose-dependent response (Figure 4D). This result, in combination with the CFD analysis, highlights the ability to locally administer drugs via the subjacent microchannel while avoiding sudden changes in the cardiac microtissues microenvironment. Moreover, the positive chronotropic response of the cardiac tissues agrees with the results from the CFD. Local dynamic drug dosing is a feature of interest for drug assays, in which the pharmacokinetics – the change of effective drug concentration over time – is an important factor to consider. ^[53]^ Moreover, the localized administration in small volumes is also important for assays that aim to study paracrine intercellular cross-talk in cardiophysiology, since having large volumes of cell culture medium between different cell types may attenuate the magnitude of the effect under study due to dilution of active chemical species.

Altogether, we were able to demonstrate the positive chronotropic response of the fabricated cardiac microtissues to isoprenaline, highlighting the physiological functionality of the tissues.

### 2.5. Co-culture of endothelial monolayers and cardiac microtissues in the Heart-on-Chip

Once regarded as an inactive epithelial tissue, the endothelium has been revealed to play crucial roles in cardiac health and disease. ^[54]^ ECs are abundant in the heart forming capillaries that irrigate the cardiac tissue ^[55]^ and are involved in several aspects of cardiac physiology, such as paracrine signalling, ^[56,57]^ fatty acid metabolism, maturation, and differentiation during development ^[58,59]^ to name a few. Moreover, the endothelium is constantly exposed to a frictional force caused by the blood flow that is important for ECs metabolism. ^[60]^ *In vitro*, ECs have been shown to express different phenotypes when cultured under shear stress, such as elongation parallel to the flow direction ^[61]^ and changes in permeability of the formed monolayer. ^[62]^

The porous membrane that separates the cardiac compartment from the microfluidic channel does not fully recapitulate on its own the fluidic path that drugs would undergo *in vivo*. *In vivo*, endothelial cells form a monolayer that line the cardiac vasculature, providing a supporting barrier to the adjacent cardiac tissue.

Aiming to mimic this configuration in the HoC, we established a co-culture with hPSC-ECs lining the subjacent channel previously used for localized drug administration of the cardiac microtissues (**Figure 5**A). We assessed the confluence of the hPSC-EC monolayer monocultured in the HoC by fluorescently labelled F-actin filaments and hPSC-ECs were able to fully cover the subjacent channel in monoculture and co-culture conditions (Figure 5B). Since the area of interaction between the two compartments is limited to the shaft of the cardiac compartment, we investigated the coverage of the hPSC-ECs at this location in co-culture conditions and confirmed that the hPSC-ECs were able to form a continuous monolayer lining the subjacent channel (Figure 5B).

**Figure 5.**
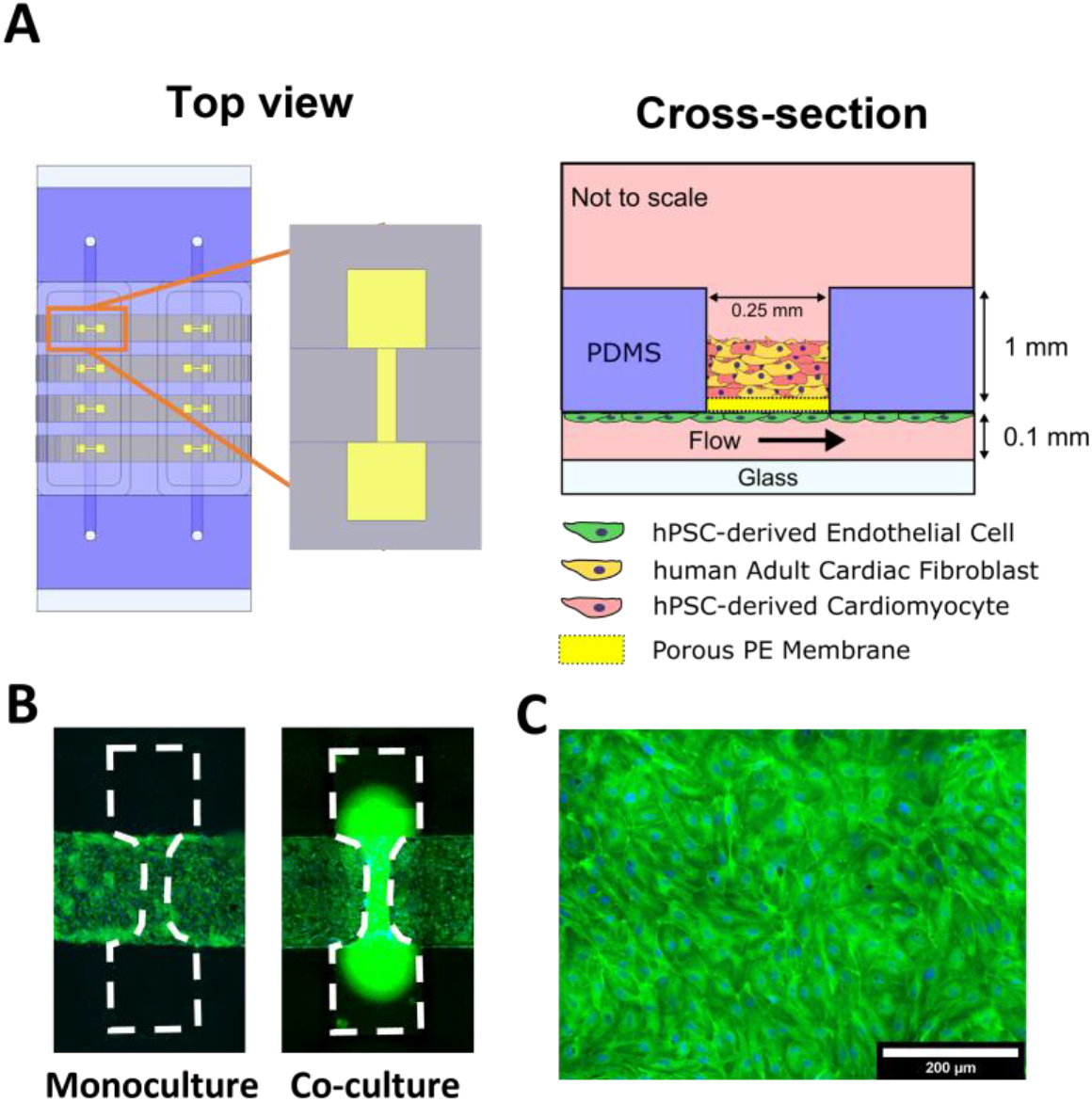
Endothelialisation of the subjacent microfluidic channel. A) On the left, schematic top view of the HoC device and the dumbbell-shape cardiac compartment with the subjacent microfluidic channel. On the right, schematic cross-sectional view of the culture compartments and its respective cell types. B) Endothelialisation of the microfluidic channel in monoculture and co-culture conditions, both demonstrating good coverage of the microfluidic channel. C) Fluorescent microscopy micrograph of a closed-up view of the hPS-ECs cultured for up to 5 days on a rocking platform without any evident cell alignment. Flow direction was from left to right and vice-versa. In both B and C, the cytoskeleton of the hPS-ECs was stained with actin-green and their nuclei with DAPI in blue.

The use of a custom-made rocking platform reduced labour-intensive setup of microfluidic pumps and ensured, via mass transport, that hPSC-ECs were cultured in a homogeneously distributed nutrient medium as opposed to the gradients that typically develop over the length of microfluidic channels in static cultures. The versatile design of our device still enables active medium perfusion using pressure-driven or syringe pumps to attain higher control and stability on fluid flow through the microfluidic chip if desired.

In our setup, the shear stress generated from the gravity driven-flow was calculated to reach a maximum of 0.44 Pa (**Supplementary Figure 7**), equivalent to a vein microenvironment. ^[63]^ Due to the high width-to-height ratio of the microchannel, the fluid flow front gets closer to a planar shape reducing the characteristic parabolic profile of the Poiseuille pressure-driven flow front (**Supplementary Figure 8**).

However, under these conditions of low shear and bi-directional flow, no evident hPSC-ECs alignment could be detected in our culture setup (Figure 5C). The low shear stress magnitude and the consideration that the mechanisms enabling endothelial cells to sense shear forces require the convergence of several different pathways, ^[60]^ may explain the lack of cell alignment to flow direction. Moreover, certain phenotypes do not align in response to shear stress ^[64]^ and immaturity of hPSC-ECs is a known shortcoming of hiPSC-derived cells that typically resemble an embryonic stage. ^[65]^ Elucidation of the factors at play in mechanical stimulation is an ongoing subject of research. Characterizing the phenotype of hPSC-ECs and evaluating their maturity will be a key factor on the recapitulation of the *in vivo* microenvironment in HoC systems like the one here presented.

We co-cultured endothelial cells with the cardiac microtissues for a maximum period of 5 days on a rocking platform until a monolayer was obtained (Figure 5B). Upon co-culture, no changes in the morphology of hPSC-ECs were detected and no significant alterations were found in the beating frequency of cardiac microtissues.

Altogether, the ability to culture a monolayer of hPSC-ECs in the microfluidic channel will enable future studies in cardiac-endothelial cross-talk.

## 3. Conclusion

We have successfully developed a HoC model that integrates hPSC-derived cardiac microtissues and demonstrated that the microtissues in the HoC can be paced at voltages that are lower than what has ever been reported due to the integration of 3D pyrolytic carbon electrodes and cell attachment to the electrodes. We also demonstrate that cardiac tissues respond to inotropic drugs that are dosed in the subjacent microfluidic channel with an increasing beat frequency trend, typical of isoprenaline. Overall, our HoC represents a unique platform for studying cardiac physiology and drug responses *in vitro* in a versatile open-top microfluidic device.

Other HoC devices that integrate cardiac microtissues with adjacent compartments have been previously reported. ^[29,66]^ Compared to these models, our design presents an improvement on the ease of fabrication without sacrificing the anatomical relevance of the model. The design of our device also follows the conventional organ-on-chip configuration of two compartments with a semi-permeable membrane. This presents advantages on the formation of a tissue-tissue interphase that more closely mimics the heart and may facilitate future integration into ‘body-on-chip’ systems in which several organs-on-chips are linked by identical vascularized compartments. ^[67]^ In comparison to the other reported HoC models, the fluorescent imaging of our cardiac microtissues showed that hPSC-CMs were more homogenously distributed, more densely compacted and had sarcomeric structures in the fabricated cardiac microtissues, which altogether better resemble the *in vivo* cell-ECM ratio of the human heart.

One unique aspect of our HoC device is the open-top configuration of our design. This allows easy access to the cardiac tissue compartments during and after culture. The controlled geometries of the 3D cardiac tissue compartments enable a high degree of control over the tissue morphology, including those that show benefits on 3D cardiac tissue formation. ^[25]^

Moreover, the open-top configuration allowed the optional integration of an E-lid for electrical stimulation of cardiac tissues.

Improvement of models like the one here presented must be accompanied by development in the field of stem cell technologies where efforts are being made to enhance the maturity and phenotype of differentiated cells, bringing them closer to an adult state and away from the current embryonic or fetal phenotypes. ^[68]^ Due to the non-standardized fabrication methods in the organ-on-chip field, throughput of HoC platforms like ours is another limiting factor where this organ-on-chip technology could benefit from further improvements. Although we aimed at lowering the threshold for performing assays in a HoC, the field is still far from being able to compete with high-throughput technologies such as high-content imaging and automated cell culture. ^[69]^ Future efforts are needed to combine our HoC with microfluidic platforms for automation of perfusion ^[70]^ and sensing. ^[71]^

Organ-on-a-chip technology presents unprecedented opportunities to closely mimic *in vitro* the dynamic microenvironment that cells are exposed to in the human body. The further integration of 3D cell culture with hPSC-derived cell technologies opens a wealth of tools to introduce human physiology in HoC models. Additionally, the high level of control over mechanical, biochemical, and electrical stimuli in these cell culture platforms ensures that HoC models better recapitulate the *in vivo* microenvironment of tissues than common cell culture hardware. This microenvironment plays an important role on providing the correct cues for these to maintain and in some cases mature their phenotype.

The development, validation and use of HoC models like the one presented here will enable the realization of a more informed drug discovery process. There, OoCs will be used to their full potential in the assessment and discovery of intricate biological questions where the use of animal models fail to replicate human physiology and current cell culture techniques lack the complexity the human body.

## 4. Experimental Section

### Chip design and Fabrication

The HoC device is comprised of two independent microchannels which are each connected to their own respective set of four cardiac compartments. The cardiac compartment and microchannels are fluidically connected through a 10 μm thick polyester (PE) porous membrane with 8 μm diameter pore-size (GVS Life Sciences, USA). The cardiac compartments consist of dumbbell-shaped wells similar to what has been reported previously ^[25]^ in which cardiac microtissues were fabricated. The cardiac dumbbell-shaped wells are 3.2 mm in their longest axis, where the two squares at the edges are 1 mm^2^ connected by a 1.2 mm by 0.25 mm shaft and a total depth of 1.5 mm. Above the cardiac tissue compartments lies the cardiac reservoir that can hold approx. 450 μL when reversibly closed by a glass seal. The microchannels have a rectangular cross-section with dimensions of 1.2 mm width, 0.1 mm height and a length of 33 mm.

Devices were made out of polydimethylsiloxane (PDMS) and fabricated using an injection molding-like technique with a pair of negative replicate molds similar to a technique previously reported. ^[72]^

The HoC and the two respective negative molds were designed using SolidWorks (Dassault Systèmes, France). A CNC machine (Datron Neo, Datron AG) was used to micromill the molds using 8 mm thick casted poly(methyl methacrylate) (PMMA) (Altuglass, France) as a stock material. Dimensions of the molds were then verified by optical microscopy using a polydimethylsiloxane (PDMS) (Sylgard 184 Silicone elastomer kit, Dow corning, USA) casted replica.

The HoC was fabricated by sandwiching 1 mm wide PE membrane strips perpendicularly aligned to the channel between the two negative molds. The membrane strips were held in place by double-sided tape (3M, USA). The two PMMA molds were brought together and clamped with two N46 neodymium nickel-plated square magnets rated with ca. 58.8 N force (Webcraft GmbH). PDMS was mixed in a 10:1 (wt:wt) polymer base to crosslinker agent ratio, degassed and injected with a 12 ml syringe (BD plastics) into the assembled mold. Air bubbles formed during PDMS injection were allowed to escape for 30 min by leaving the mold in a vertical position at room temperature (RT) on the lab bench followed by a 65°C overnight incubation in an oven for PDMS cross-linking. After curing, molds were disassembled, the PDMS devices were peeled off from the molds and edges were trimmed with a scalpel. Inlets for the microchannels were opened with a 1 mm diameter biopsy puncher (Robbins Instruments, USA).

Glass coverslips with dimensions 50×24×0.15 mm (Menzel Glazer, Thermo scientific, USA) were spin-coated (1500 rpm, 30 s, 1000 rpm/s, Spin150, Polos, The Netherlands) with PDMS and cured overnight. HoC and PDMS-coated coverslips were simultaneously exposed to air plasma (50 W) for 40 s (Cute, Femto Science, South Korea), bonded and incubated at 65°C for at least 3 h to enhance bonding.

### Chemical chip surface functionalization

Fully assembled devices were silanized to enhance hPSC-ECs adhesion to the microchannels. Devices were first exposed to air plasma (50 W) followed by filling channels with an aqueous solution of (3-Aminopropyl)triethoxysilane (APTES) (Sigma Aldrich, Germany). Devices were then incubated for 1 min at RT, submerged in 100% ethanol and channels flushed with pure ethanol. Devices were then air blown dried with nitrogen and incubated in an oven at 65°C to complete ethanol evaporation. After incubation, UV-sterilization was performed during 30 min in a laminar air flow cabinet (Telstar, The Netherlands). Microchannels were coated with a rat tail collagen type I (VWR) solution of 0.1 mg/ml in DPBS (rat tail collagen I, ThermoFisher, USA), and incubated for 30 min at 37°C in a humidified incubator. The collagen solution was then flushed out of the channels, after which they were filled with DPBS until cell seeding.

### 3D pyrolytic carbon electrode fabrication

For electrical pacing of the cardiac microtissues, the glass seal was substituted by an electronic lid, ‘E-lid’, containing carbon electrodes. The carbon electrodes were fabricated using a combination of common photolithography technology and 3D printing in a three-step fabrication process, namely photolithography for patterning of connection pads and tracks, 3D printing of the pillars and pyrolysis to convert the polymer precursor structures into pyrolytic carbon.

First, the connection pads, leads and base of the E-lid were fabricated using photolithography. Briefly, a 600 nm layer of SiO_2_ was grown on a 4-inch silicon wafer by thermal oxidation. The wafer was then dehydrated for 24 h at 250°C and plasma activated for 2 min at 500 W and 120 sccm of oxygen (300 Plasma Processor, PVA TePla, Germany). A 15 μm thick SU-8 layer (SU-8 2035, Kayaku Advanced Materials, USA) was then spin-coated onto the surface, followed by a soft bake step for 1 h at 50°C. The SU-8 film was exposed to UV light to define the pattern of the 2D electrode precursor structures using a maskless aligner (MLA 100 Maskless Aligner, Heidelberg Instruments, Germany) at a dose of 250 mJ/cm^2^ with a wavelength of 365 nm. This was followed by a post-exposure bake for 1 h at 50°C to crosslink the SU-8. The wafer was then developed using Propylene glycol monomethyl ether acetate (PGMEA, mr-Dev 600, micro resist technology GmbH, Germany) removing the un-crosslinked SU-8. The SU-8 structures were then flood exposed (MA6/BA6 mask aligner, Süss MicroTec, Germany) with four exposure cycles at 250 mJ/cm^2^ each with 30 s intervals between exposures. A final hard bake step was done on a hotplate for 15 hours at 90°C to increase the degree of crosslinking of the SU-8. The thickness of the patterned photoresist was verified by profilometry (DektakXT, Bruker, USA) and the wafer was subsequently diced (DAD 321, DISCO, Japan) into individual E-lids, each wafer containing 3 chips.

The 3D printed pillars were designed with heights ranging from 3 to 5 mm and diameters ranging from 300 to 800 μm using CAD software (Fusion 360, Autodesk, USA) and 3D printed with a stereolithography 3D printer (Form 2, Formlabs, USA) using a photosensitive resin (High Temp Resin V1, Formlabs, USA). The selected design had a height of 4 mm and a diameter of 600 μm. First, to align the 3D printing of the pillars to the diced E-lids, a holder structure was printed and washed with isopropyl alcohol (IPA) for 15 min (Form Wash, Formlabs, USA). The diced E-lids were then activated in an O_2_ plasma at 120 W for 75 s (Zepto, Diener electronic, Germany), mounted onto the previously printed holder using double-sided tape and pillars were then printed onto the chips. This was followed by a washing step with IPA for 15 min and a curing step of 1 h at 60°C (Form Cure, Formlabs, USA). The height of the 3D printed pillars was 3.62 mm and the diameter was 432 μm, which is slightly lower than the design values.

Lastly, the final E-lids with the 3D printed pillars were pyrolyzed achieving conversion of the precursor polymers to pyrolytic carbon. The chips were loaded into a high temperature tube furnace (OTF-1200X, MTI Corporation, USA), a constant flow of nitrogen gas of 1000 sccm was applied and pyrolysis was conducted in three temperature steps. First, the temperature was ramped up to 375°C at 10°C/min and maintained for 3 h, then ramped up to 425°C at 12°C/min and maintained for 30 min. and lastly ramped to 900°C at 10 °C/min and maintained for 1h for annealing and cooled down at 10 °C/min. The final height of the pyrolytic carbon micropillars was 1.45 mm and the diameter was 158 μm, which demonstrates the considerable shrinkage compared to the 3D printed pillars due to mass loss during pyrolysis.

### Conductivity measurement and cyclic voltammetry

To verify the electrical conductivity of the carbon leads and the carbon micropillars of the E-lid and that these were electrically connected to the underlying 2D electrodes, the electrical resistance between the two sets of 4 carbon micropillars was measured using carbon paste and a multimeter.

Cyclic voltammetry was used to identify if the 3D pillar electrodes on the E-lid were electroactive. The electrode terminals were connected to a potentiostat (PalmSense4, PalmSens, The Netherlands) using crocodile clips (AGF1, Hirschmann, The Netherlands). The E-lid was used as a working electrode along with a Ag/AgCl 3 M KCl3MKCl reference electrode (PalmSens, The Netherlands) and a Pt counter electrode in an equimolar solution of 10 mM ferri/ferrocyanide in phosphate buffered saline (PBS) at a pH of 7.4. A scan rate of 10 mV/s was used for all measurements.

### Cell differentiation and culture

Cardiomyocytes (CMs) were derived from an embryonic human pluripotent stem cell line with two reporter genes for the transcription factor NKX-2.5GFP/+ and the fluorescently tagged sarcomere accessory protein α-actinin with GFP and mRuby, respectively and differentiated as previously reported. [37] Briefly, hESCs were seeded at a density of 25×10^3^ cell/cm^2^ on Matrigel-coated 6-well plates in Essential 8 medium (ThermoFisher, USA) on day −1. At day 0, mesodermal differentiation was initiated by addition of Wnt activator CHIR99021 (1.5 μmol/L, Axon Medchem 1386), Activin-A (20 ng/mL, Miltenyi 130–115-010) and BMP4 (20 ng/mL, R&D systems 314-BP/CF) in Bovine Serum Albumin (BSA) Polyvinylalcohol Essential Lipids (BPEL) medium. At day 3, Wnt was inactivated by adding XAV939 (5 μmol/L, R&D Systems 3748) in BPEL. ^[73]^ Cell cultures were refreshed with BPEL on day 7 and 10 after the start of differentiation until differentiation was completed. CMs were then frozen in medium C containing 50% Knock out serum, 40% BPEL, 10% DMSO and 1x RevitaCell and stored in liquid nitrogen. Before CMs were thawed, 6-well well-plates were first coated with vitronectin (5 μg/mL, Thermo Fisher, USA) followed by a coating with 10% fetal bovine serum (ThermoFisher, USA) in DMEM (Sigma, USA) for 1 h and 30 min at 37°C in a humidified incubator. CMs were thawed and plated on coated 6-well plates at a cell density of 1×10^5^ cell/cm^2^ and cultured in cardiomyocyte maturation medium ^[74]^ supplemented with 100 nM triiodothyronine hormone (T3) (Sigma-Aldrich), 1 μM dexamethasone (Tocris) and 10 nM LONG R3 IGF-1 (IGF, Sigma-Aldrich, USA) (CM+TDI) for 3 to 4 days (16-17 after differentiation) prior to the 3D tissue fabrication.

hPSC-ECs were differentiated from hiPSC as previously reported. ^[75]^ Briefly, hiPSCs were maintained in TeSR™-E8™ medium (StemCell Technologies, Canada) on vitronectin-coated 6-well plates and seeded at day −1. 24 h after seeding, E8 medium was replaced with BPEL medium supplemented with 8 μM CHIR. On day 3, the medium was replaced with BPEL medium supplemented with 50 ng/ml VEGF (R&D systems) and 10 μM SB431542 (Tocris Bioscience) and refreshed on days 6–9. hiPSC-ECs were isolated on day 10 using CD31-DynabeadsTM (Invitrogen) as previously described. ^[75]^ hPSC-ECs were cryo-preserved after isolation and obtained batches were used in all further experiments. After thawing, hPSC-ECs were subcultured and expanded in endothelial serum free medium (SFM, ThermoFisher, USA) supplemented with 30 ng/ml VEGF (R&D Systems), 20 ng/mL bFGF (Milteny Biotech) and 1% platelet-poor plasma-derived serum (BioQuote). ^[75]^ Cells were sub-cultured on 0.1 mg/ml rat tail collagen type I T-75 coated flasks and passaged at 80% confluency. Cells were used at P2.

Human adult cardiac fibroblasts (cFBs, Bio-Connect) were subcultured following manufacturer’s instructions in T-75 culture flasks with FGM-3 medium (Bio-Connect). cFBs were used between P4 and P7.

CMs, cFBs and ECs were washed with DPBS and dissociated with 1× TrypLE select (ThermoFisher, USA) for 3 min. in a humidified incubator at 37°C. TrypLE was diluted with DMEM supplemented with FBS for CMs, FGM-3 for cFBs and supplemented SFM for ECs. Cells were then centrifuged at 240 ×g for 3 min and the supernatant was aspirated. The percentage of CMs in the resulting differentiated population was quantified by flow cytometry (MACSQuant VYB flow cytometer, Miltenyi Biotech) using the fluorescent reporter markers expressed by these cells.

Cardiac tissues were formed in the microfluidic chip by seeding CMs and cFBs inside a fibrin hydrogel in the microcompartments. After dissociation, CMs and cFBs pellets were resuspended at a concentration of 5.8×10^7^ cells/ml in CM+TDI supplemented with horse serum (CM+TDI+HS) and FGM-3, respectively. Thrombin from bovine plasma (ThermoFisher, USA) was mixed into the cell suspension at 1:300 ratio with a final gel concentration of 0.67 U/ml. Both cell types were added in an ice bath to a hydrogel mix with the following composition: fibrinogen from bovine plasma (2 mg/ml, Sigma-Aldrich); Matrigel (1:10 V/V, Corning); 2 times concentrated CM (2x CM); and aprotinin from bovine lung (1:150, Sigma). The final cell gel concentration was 4.0×10^7^ cells/ml and the CMs to cFBs ratio was 1:10.

4 ml of pre-polymerized gel were pipetted onto each dumbbell shape and incubated at RT for 10 min to allow hydrogel polymerization. CM+TDI+HS was then added to the cardiac compartment, and 3D tissues were cultured at 37°C and 5% CO_2_ in a humidified incubator. Medium was changed every 3 days.

Endothelial cells were cultured in the microfluidic chip by seeding a highly concentrated cell suspension followed by continuous medium refreshment. After dissociation, ECs pellets were resuspended at a concentration of 5.0×10^6^ cells/ml and seeded in the collagen-coated channels of the HoC. The device was flipped upside-down and incubated for 30 min in a humidified incubator at 37°C to promote cell attachment on the ceiling of the microchannel. After incubation, non-attached cells were removed by flushing with fresh medium. Pipette tips containing 100 μl of medium were mounted on the inlet and outlet of each endothelial channel. The seeded chips were placed on a custom-made rocking platform with a 35° tilting angle with a 30 s cycle in a humidified incubator at 37°C for 3 days until confluence was reached.

### Fluorescence and time-lapse microscopy

Nuclei and the cytoskeleton of ECs were stained to assess cell confluency in the channels as well as cell morphology. Briefly, cells were washed with PBS and fixed with 4% formaldehyde (ThermoFisher, USA) in PBS for 15 min at RT. Cells were then permeabilized with 0.1% Triton-X-100 (Sigma-Aldrich) for 5 min at RT and washed. ECs were then incubated at RT for 30 min. with the staining solution containing 2 drops/ml F-actin binding ActinGreen (ThermoFisher, USA) and 1.25 μg/ml 4’, 6-Diamino-2-Phenylindole (DAPI, ThermoFisher, USA). The staining solution was then washed with PBS.

EVOS FL Cell Imaging System and EVOS M7000 (Life Technologies) were used to image ECs and the cardiac tissues. For analysis of beating cardiac tissues, time-lapse images were captured using a Nikon EclipseTE2000-U fluorescence microscope (Nikon) with a high-speed camera (DMK 23UX172, The Imaging Source Europe GmbH, Germany). Cardiac tissue contraction analysis was performed using Openheartware. ^[26]^

### Electrical Pacing of the cardiac microtissues

Three-dimensional cardiac tissues were subjected to electrical pacing using a stimulus generator (STG4008-16mA, Multi Channel Systems MCS GmbH). Biphasic electrical pacing was performed at 1 Hz and 2 Hz at 1.6 V/cm with a stimulus duration of 20 ms in CM+TDI+HS medium.

### Treatment of Heart-on-Chip with drug

The drug was infused through the microfluidic channel subjacent to the cardiac microtissues in the HoC. All cardiac tissues were cultured for 14 days prior to the drug assay.

Isoproterenol (Sigma-Aldrich), a positive inotropic agent, was first diluted in Dulbecco’s PBS (DPBS) at a stock concentration of 10 mM. Isoprenaline administration to the cardiac tissues was performed at concentrations ranging between 1 nM and 30 nM in CM+TDI+HS medium in increasing concentrations and incubated for 5 min. before imaging.

Two pipette tips were fitted into the HoC inlets with 100 μl of the spiked medium and the device was incubated in a humidified incubator at 37°C on a rocking platform for 5 min. to ensure mass transport along the microfluidic channel. Video clips of 10 s were made for each tissue and all measurements took place within 10 min after incubation.

### Drug diffusion simulation

To gain insight into the mechanism of drug diffusion between the different compartments, computer fluid dynamic simulations were employed using COMSOL Multiphysics (COMSOL, Sweden). The simulation domain was comprised of one dumbbell-shaped microwell, the overhead volume of the reservoir and the membrane.

The latter was simulated as a custom porous material with a thickness of 10 μm and a calculated porosity of 5%, derived from the manufacturer’s specifications. The transport of diluted species module was employed and the diffusion constant for isoprenaline was considered as 10^−5^ cm^2^/s. ^[76]^ A time-dependent study was then performed with a 0.1 s interval step.

### Statistics

Results are displayed as mean ± standard error of the mean (SEM) except when stated otherwise. All analysis was performed using GraphPad Prism 8.0 (GraphPad Software, USA).

## Acknowledgements

A.V, A.v.d.B, R.P and A.D.v.d.M acknowledge the European Research Council (ERC) under the Advanced Grant ‘VESCEL’ Program (Grant no.669768) of Prof. Van den Berg. J.Y.P. and S.S.K. acknowledge funding by the European Research Council under the Horizon 2020 framework programme grant no. 772370-PHOENEEX.

