## Supplementary figures for "Generation and Culture of Cardiac Microtissues in a Microfluidic Chip with a Reversible Open Top Enables Electrical Pacing, Dynamic Drug Dosing and Endothelial Cell Co-Culture"

### Supplementary figure 1. Reversibly closed cardiac compartment

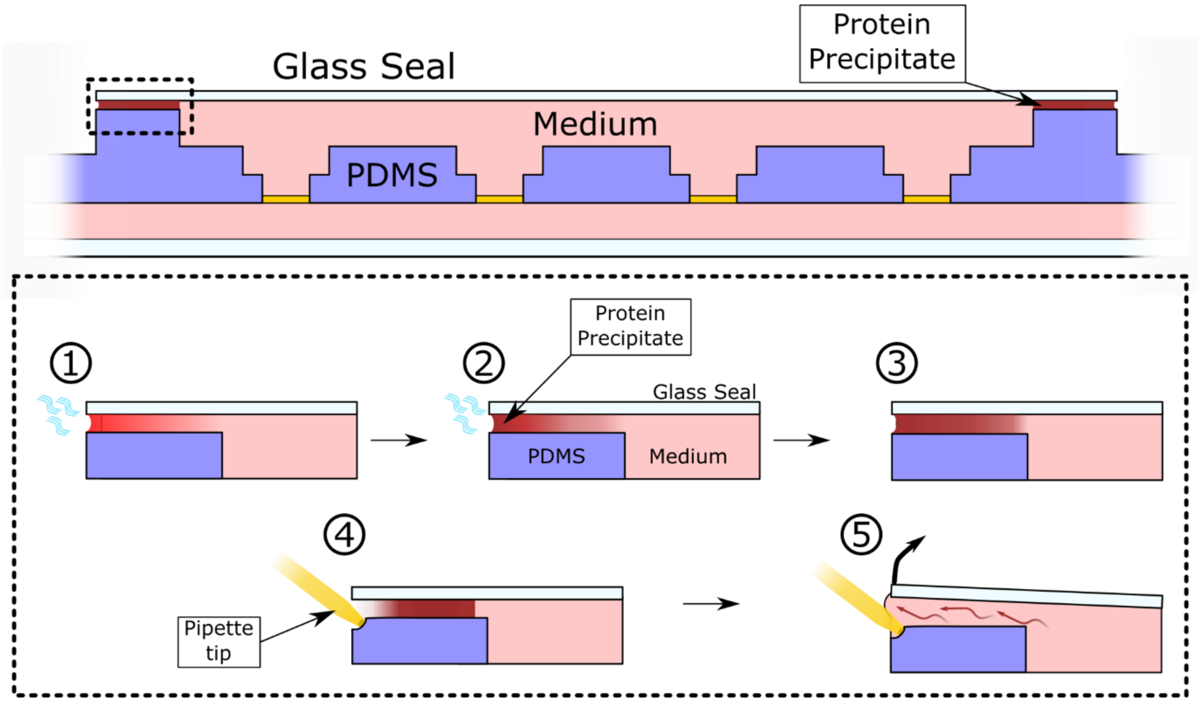


*Supplementary Figure 1. Schematic longitudinal cross-section overview of the glue-free reversible sealing method. 1) A thin film of fluid is created via capillary action between the glass seal and the designed rim enclosing the cardiac compartment; after cell seeding and incubation, water in the medium evaporates. 2) Medium water evaporation locally increases protein and salts concentrations. 3) After approximately 8h in incubation a seal is created by the protein precipitate. 4) To reopen the cardiac compartment, a pipette tip, in yellow, is used to deform the PDMS rim and disrupt the protein precipitate. 5) By capillary action medium reflows into the gap between the glass seal and the PDMS rim.*

### Supplementary figure 2. Flow cytometry and Cardiac differentiation efficiency

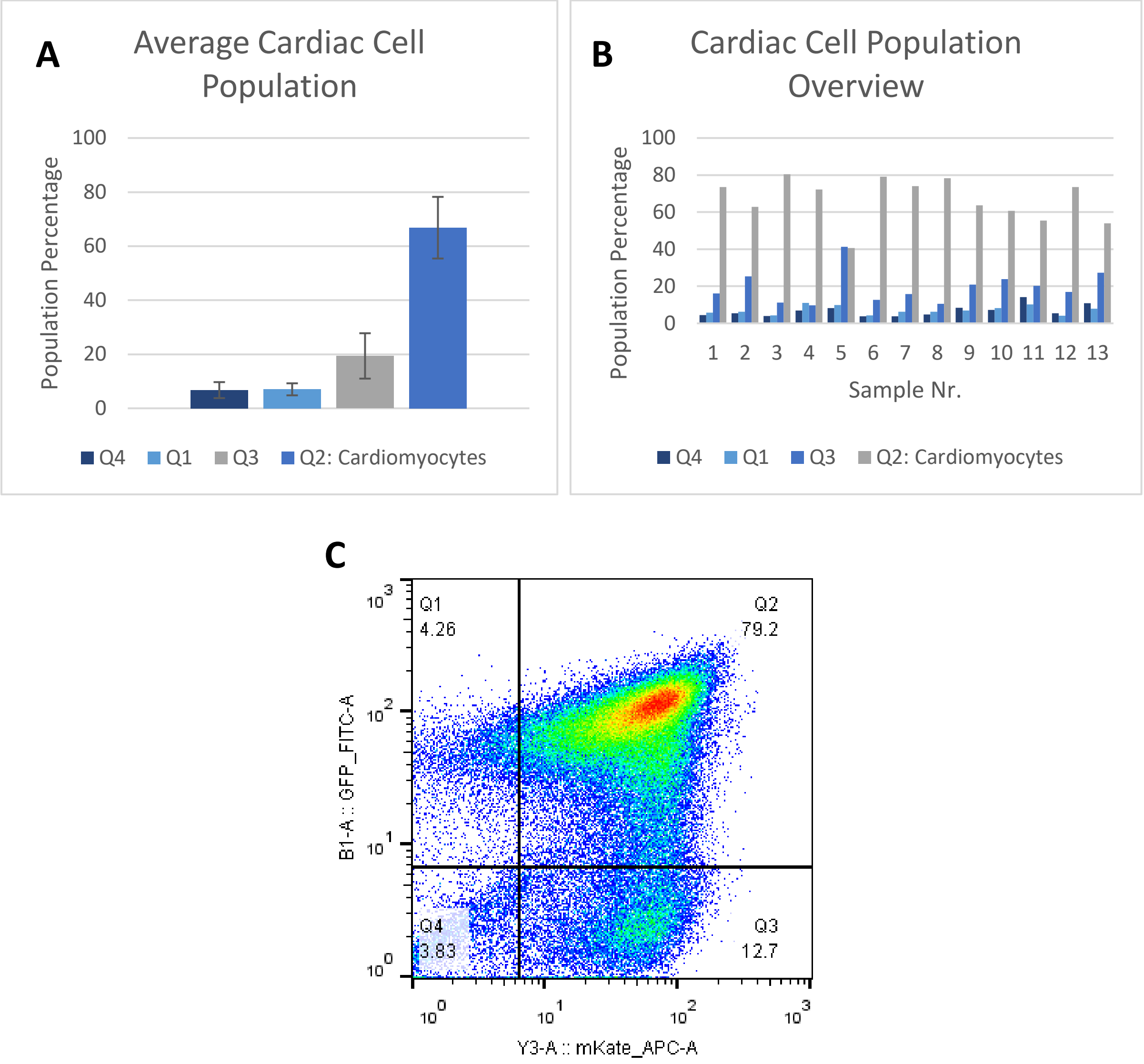


*Supplementary Figure 2. Overview of the cardiac cell population obtained after differentiation characterized using flow cytometry and the reporter genes expressed by the DRAGGN-line. A)The average cardiac cell population used to fabricate the cardiac tissues had on average the presented distribution. B) Distribution of cardiac cells in each of the cell populations used in the study. C) Flow cytometry chart depicting the different populations and respective gates. The y axis is for GFP-NKX-2.5 and x for mRubyII-α-actinin.*

#### Supplementary figure 3. Addition of cFBs and tissue remodeling tissue compaction and out of the shape

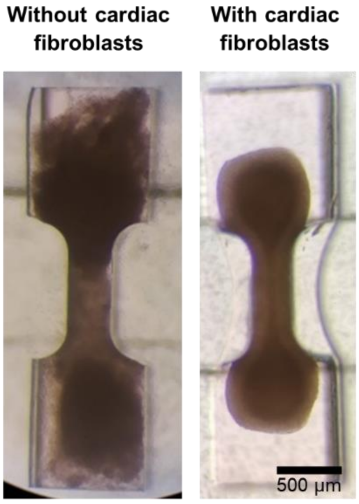


*Supplementary Figure 3. Cardiac tissue compaction without and with human adult cardiac fibroblasts. Left – Depicts the cardiac tissue without cFBs after three days in culture with poor defined boundaries and apparent non-homogeneous cells distribution. Right – depicts the cardiac tissue in co-culture with cFBs after three days in culture with well-defined tissue boundaries and homogeneous cell distribution.*

#### Supplementary figure 4. Dimension differences among the different designed, 3D printed and pyrolyzed micropillar electrodes fabricated.

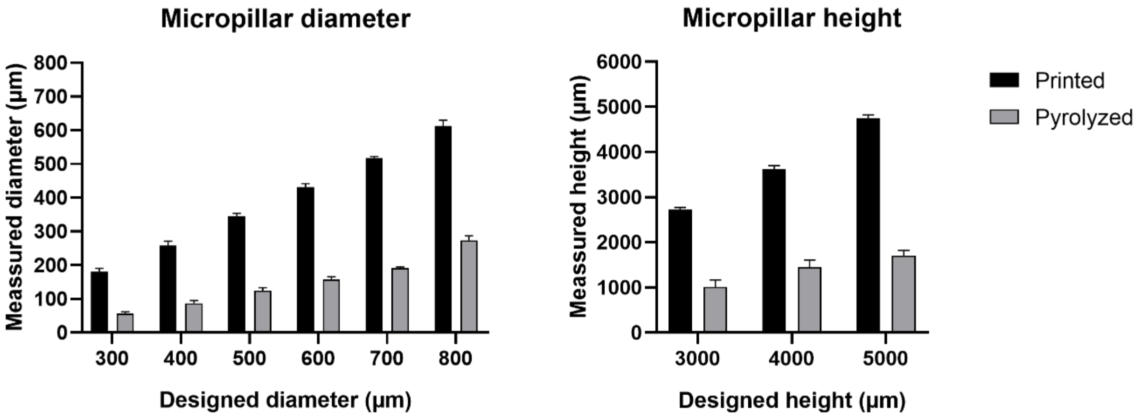


*Supplementary Figure 4. Dimension differences among the different designed, 3D printed and pyrolyzed micropillar electrodes fabricated. The different micropillars designs varied the diameter, ranging from 300 to 800 µm, and height m ranging from 3mm to 5 mm. The designs were 3D printed, pyrolyzed and measured at each step of the process for selecting the best fitting design dimensions. Error bars represent standard deviation.*

#### Supplementary figure 5. Excitation threshold detection using Fourier transform and a 2 Hz stimulation frequency.

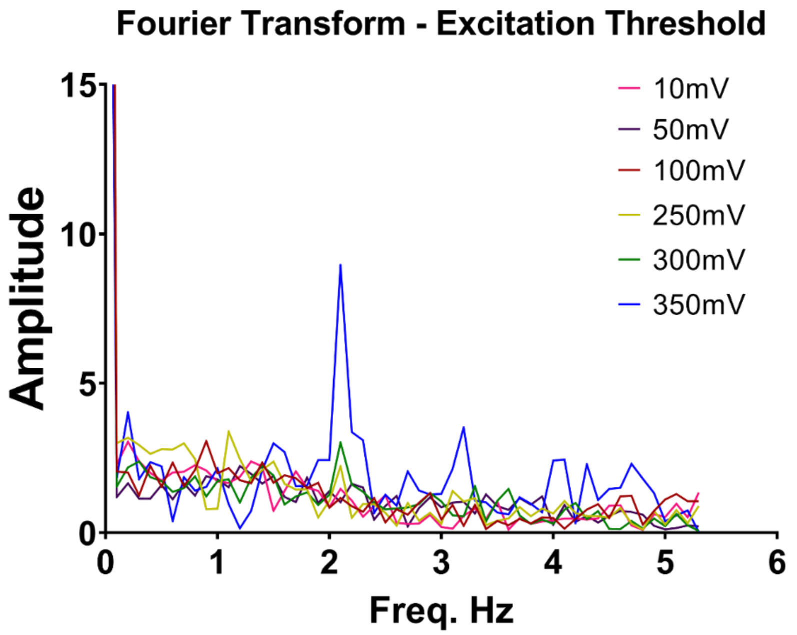


*Supplementary Figure 5. Fourier transform derived from the beating frequency of the cardiac tissue and measured with kymography. Trains of bi-phasic, square-shaped electric pulses with 20ms duration and 2 Hz frequency were applied using the E-lid with increasing amplitudes ranging from 10 to 350 mV. The excitation threshold was determined by the appearance of the electric excitation frequency in the Fourier transform of the beating signal of the cardiac tissue.*

#### Supplementary figure 6. Summary graph of the individual response of each of the 3D cardiac tissues to the different isoprenaline concentrations tested.

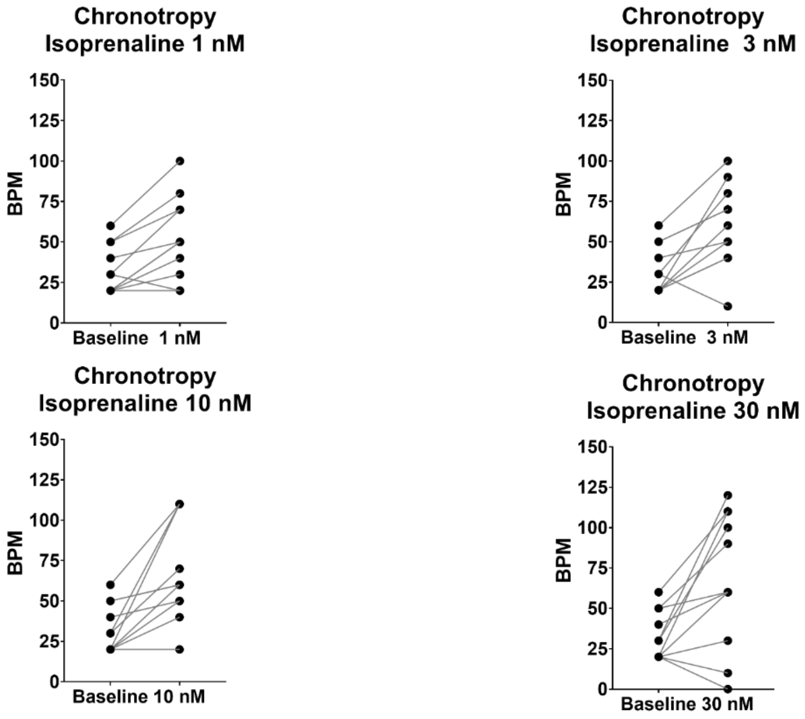


*Supplementary Figure 6. Summary graph of the individual response of each of the 3D cardiac tissues to the different isoprenaline concentrations tested.*

### Supplementary figure 7. Fluid flow shear stress on rocking platform

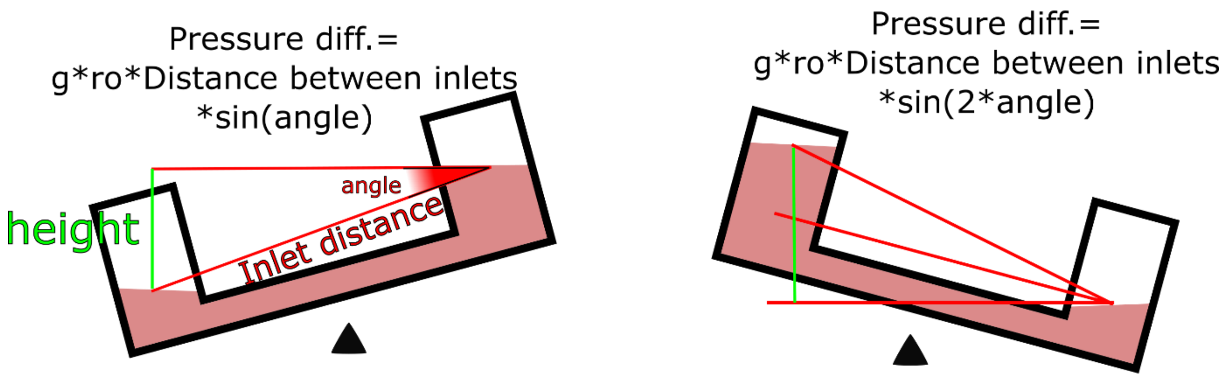


$$\alpha = 35°$$

$$\sin 2\alpha = \frac{\Delta h}{d} \Leftrightarrow \Delta h = 31\, mm$$

$$\Delta P = \Delta h \rho g \Leftrightarrow \Delta P = 303{,}9\ Pa$$

$$R = \frac{12\mu L}{wh^3} \Leftrightarrow R = 2{,}4 \times 10^{11}\ Pa.s^{-1}.m^{-3}$$

$$\Delta P = R \times Q \Leftrightarrow Q = 1{,}26 \times 10^{-9}\ m^3.s^{-1} \Leftrightarrow Q = 75{,}9\ \mu L.min^{-1}$$

$$\tau_{peak} = \frac{6\mu Q}{wh^2} \Leftrightarrow \tau = 0{,}44\ Pa$$

*Supplementary Figure 7. Schematic representation of the gravity-driven flow stablished with a custom-built rocking platform and resulting shear stress calculation. Top left, in the first cycle, the height difference can be derived from trigonometry relation where the tilting angle is that of the rocking platform. Top right represents the second and subsequent rocking cycles where the same relations are derived but this time with double the angle of the rocking platform due to the volume imbalance between the two inlets.*

### Supplementary figure 8. Homogeneous shear rate in vascular compartment.

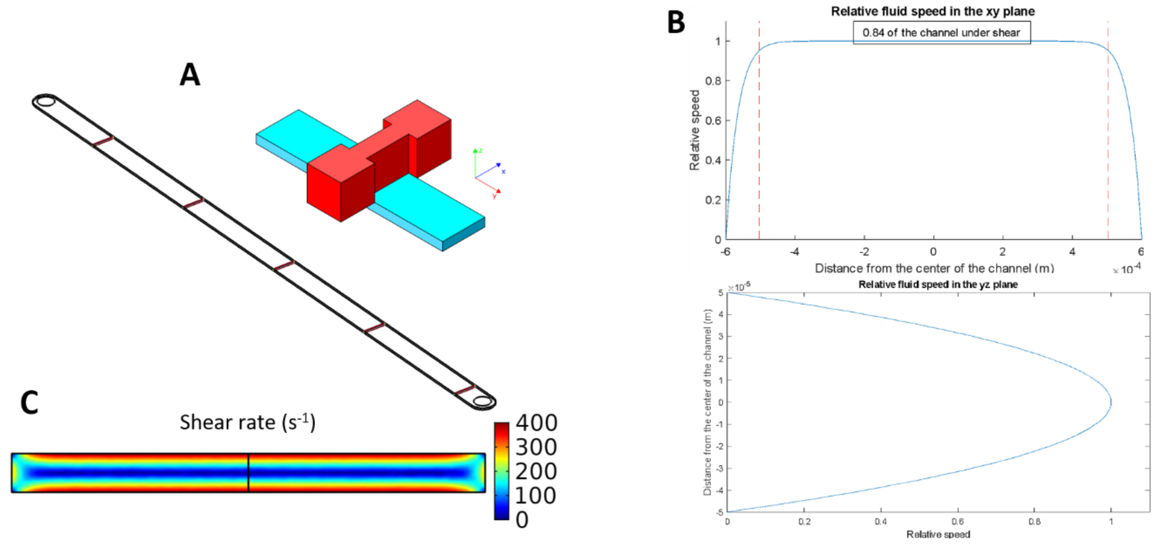


*Supplementary Figure 8. Shear rate homogeneity and magnitude experienced by endothelial cells in the vascular compartment. A) Schematic isometric view of the culture compartments, cardiac compartment in red and vascular compartment in turquoise and the isometric view of the vascular compartment used in the simulation. B) Flow profile in the vascular compartment in a top and side view. C) Computer fluid dynamics modelling of the shear rate experienced by the cells in a front view – XZ plane.*
